# Thermorheological mapping and molecular insights into salt-dependent gel-like network formation in *Mortierella alpina* chitin-like exopolysaccharide solutions

**DOI:** 10.64898/2026.09.05.749623

**Authors:** Luis Daniel Goyzueta-Mamani, Haruna Barazorda-Ccahuana, Miguel Daniel Noseda, Rilton Alves de Freitas, Carlos Ricardo Soccol, Júlio Cesar de Carvalho

**Author notes:** Corresponding author: Luis Daniel Goyzueta-Mamani.

## Abstract

Fungal chitin-like exopolysaccharides are promising fermentation-derived materials, however the molecular-level determinants of their salt- and temperature-dependent gel-like behavior remain poorly understood. Here, a previously characterized *Mortierella alpina* exopolysaccharide was re-examined by integrating oscillatory rheology with all-atom molecular dynamics simulations. Frequency sweeps across 5-15 mg mL^−1^, two ionic media (0.154 mol L^−1^ NaCl and 0.28 mol L^−1^ LiCl), and 10-60 °C were used to map liquid-like, transitional, and elastic-dominated regimes. Increasing concentration strengthened network behavior, but the two salts followed distinct pathways: NaCl promoted a more coherent progression toward elastic dominance and more thermorheologically compatible relaxation behavior, whereas LiCl produced a high-dissipation intermediate regime and stronger temperature dependence. Simulations of a minimal multichain model comprising eight GlcNAc10 oligomers at approximately 15.8 mg mL^−1^ showed that NaCl favored larger, more compact assemblies despite weak and short-lived Na^+^-oxygen coordination. LiCl formed sharper, longer-lived first-shell contacts but maintained smaller and more expanded assemblies. Ion-mediated interchain bridges were sparse and transient. The combined results show that local cation binding strength does not directly predict collective network formation and support a dynamic physical-network mechanism governed by salt-specific coupling among hydration, ion exchange, chain packing, and reversible interchain association.

## 1. Introduction

Chitin is a linear (1→4)-linked poly(β-*N*-acetyl-D-glucosamine) and one of the most abundant structural polysaccharides in nature (Pillai et al., 2009; Rinaudo, 2006). Most technological development has focused on crustacean shells and on chitosan, because native chitin is highly crystalline and difficult to process in conventional aqueous media (Kaur & Dhillon, 2015; Pillai et al., 2009; Synowiecki & Al-Khateeb, 2003; Younes & Rinaudo, 2015). Fungal production offers a complementary route in which biosynthesis, composition, and downstream handling can be controlled under fermentation conditions (Steinfeld et al., 2019; Teng et al., 2001). Extracellular fungal chitin-like polymers are particularly interesting because they may combine a highly acetylated backbone with solution processability that differs from conventional shell-derived chitin.

The *Mortierella alpina* EPS investigated here was previously characterized as a highly acetylated GlcNAc-rich chitin-like polysaccharide, composed predominantly of (1→4)-linked poly(β-*N*-acetyl-D-glucosamine) residues, with a weight-average molar mass of approximately 4.9 × 10^5^ g mol^−1^ and a degree of acetylation above 90% (Goyzueta M. et al., 2020). The original study established its solubility in concentrated LiCl and NaCl media and documented concentration- and temperature-sensitive viscoelastic behavior. However, those data were interpreted mainly phenomenologically, and the relationship between salt-specific local coordination and collective network formation remained unresolved.

Oscillatory rheology is well suited to this problem because storage modulus (G′), loss modulus (G″), complex viscosity (|η*|), and loss tangent (tanδ = G″/G′) report on different aspects of stress storage, dissipation, and relaxation (Chambon & Winter, 1987; Picout & Ross-Murphy, 2003; Stojkov et al., 2021; Winter & Chambon, 1986; Winter & Mours, 1999). Near a critical gel, the Winter-Chambon framework predicts similar power-law exponents for G′ and G″ and a frequency-independent phase angle (Chambon & Winter, 1987; Winter & Chambon, 1986). Physical biopolymer networks often depart from this ideal behavior, but tanδ maps, crossover analysis, concentration scaling, and time-temperature superposition remain useful for distinguishing simple thermal shifting from changes in network topology (Chen et al., 2026; Picout & Ross-Murphy, 2003; Pieklarz et al., 2022; Zhao et al., 2003).

The response of highly acetylated amino polysaccharides to ionic media cannot be inferred from cation charge alone. Ionic strength and acetylation influence chain dimensions and association, whereas Li^+^ and Na^+^ differ markedly in hydration-shell organization and exchange dynamics. Recent experimental and simulation studies further indicate that chitin self-assembly results from a balance among hydration, hydrogen bonding, hydrophobic association, and interchain packing (Mähler & Persson, 2011; Romany et al., 2023). These observations suggest that stronger local cation–oxygen coordination, predominant in highly acetylated GlcNAc-rich chains, may not necessarily promote more extensive collective aggregation (Narkevicius et al., 2019; Rinaudo et al., 1993). Recent polysaccharide studies have similarly combined rheology and atomistic simulation to distinguish local chain interactions from emergent network assembly (McGregor et al., 2025).

We hypothesized that NaCl and LiCl would not act as interchangeable monovalent media, because differences in cation hydration and polymer-oxygen coordination should alter the coupling between local ionic interactions and multichain association. The objective of this research was therefore to integrate quantitative thermorheological mapping with all-atom molecular dynamics simulations of GlcNAc10 oligomers. This combined approach was designed to determine whether strong local ion binding promotes, hinders, or is decoupled from the emergence of a collective gel-like network.

While the previous report described the overall concentration- and temperature-dependent viscoelastic behavior, the present study derives previously unreported tanδ state maps, critical-gel diagnostics, concentration-scaling descriptors, time–temperature superposition behavior, and atomistic ion-coordination mechanisms from the complete dataset. Beyond this specific fungal EPS, the study tests a broader physicochemical principle relevant to hydrated polysaccharide networks: stronger local cation coordination does not necessarily translate into stronger collective polymer association or macroscopic network reinforcement. This mechanistic framework may support the rational formulation and processing of chitin-like EPS systems.

## 2. Materials and methods

### 2.1 EPS identity and rheological dataset

This study reanalyzed the rheological dataset of the chitin-like EPS produced by *M. alpina* under submerged fermentation and previously characterized by (Goyzueta M. et al., 2020). The established structural descriptors are summarized in Table 1.

**Table 1.** Structural characteristics and experimental design of the fungal chitin-like EPS. Structural descriptors from (Goyzueta M. et al., 2020); experimental design and rheological formulations established for the present study.

| Characteristic | Value / condition | Source or method |
| --- | --- | --- |
| Polymer identity | Highly acetylated GlcNAc-rich chitin-like EPS, predominantly composed of β-(1→4)-linked *N*-acetyl-D-glucosamine residues | Monosaccharide/linkage analysis |
| Weight-average molecular mass | $4.9 \times 10^5 \text{ g mol}^{-1}$ | Size-exclusion chromatography, single symmetric HPSEC-MALLS-RID peak |
| Degree of acetylation | >90% | $^{13}\text{C}$ CP-MAS NMR |
| EPS concentrations | 5, 10, and 15 mg mL$^{-1}$ | Rheological formulations |
| Ionic media | 0.154 mol L$^{-1}$ NaCl; 0.28 mol L$^{-1}$ LiCl | Rheological formulations |
| Temperature range | 10-60 $^{\circ}\text{C}$; LiCl 15 mg mL$^{-1}$ retained to 50 $^{\circ}\text{C}$ | Frequency sweeps |
| Experimental replication | Three replicate measurements per formulation | Mean ± SD at matched conditions |

EPS solutions were prepared at 5, 10, and 15 mg mL−1 in 0.154 mol L−1 NaCl or 0.28 mol L−1 LiCl. The lyophilized EPS was directly reconstituted in the corresponding NaCl or LiCl solutions without deliberate pH adjustment. The resulting solutions were near neutral pH. Small-amplitude oscillatory shear measurements were acquired with a Thermo Scientific Haake Rheostress 1 rheometer using a cone-and-plate geometry (40 mm diameter, 2° cone angle, 1 mm gap). Frequency sweeps were acquired from approximately 0.01 to 46.42 Hz over an experimental temperature window of approximately 10–60 °C, with the available temperature range varying among formulations. For quantitative analysis, the frequency range was restricted to ≤10 Hz to minimize potential high-frequency inertial effects associated with the measuring geometry. The imposed stress was selected within the linear viscoelastic range (0.1 Pa unless otherwise documented), and mineral oil was used to reduce evaporation.

Measurements were performed in triplicate for each salt-concentration condition. The three runs were retained separately during preprocessing and were summarized only after alignment of the temperature and frequency grids. The complete experimental matrix and verified replicate structure are provided in Table S1. Values at a fixed salt × concentration × temperature × frequency condition are reported as mean ± standard deviation (n = 3). Temperature and frequency points within the same sweep were treated as repeated observations, not as independent replicates. The 15 mg mL^−1^ LiCl data above 54 °C were excluded because of an abrupt, physically inconsistent increase in G′ and erratic tanδ values attributable to an instrumental or sample-contact anomaly.

### 2.2. Data preprocessing and rheological descriptors

Raw RheoWin and GraphPad files were consolidated into a master table containing sample identity, salt, EPS concentration, nominal and measured temperature, frequency, angular frequency, G′, G″, |η*|, strain amplitude, replicate number, and source file. Decimal separators and units were standardized. Frequency in hertz was converted to angular frequency as ω = 2πf for calculation of complex viscosity and rheological relations expressed in terms of ω.

Data quality was assessed before calculating derived descriptors. Missing or non-positive frequency values, negative moduli, non-physical viscosity values, large temperature deviations, and isolated discontinuities at the limits of the instrument window were flagged. Candidate outliers were evaluated using median-absolute-deviation screening and residual inspection; exclusion required consistency with a documented instrumental artifact. No smoothing was applied to primary calculations. Interpolation was restricted to crossover estimation and graphical maps in log-frequency space.

The loss tangent was calculated as tanδ = G″/G′, the magnitude of the complex modulus as *G* =* √*[(G′)*^2^ *+ (G″)*^2^], and the complex viscosity as |*η\**| *= G* / ω*. Crossover frequencies were estimated where log(*G*′) − log(*G*″) changed sign. Local scaling exponents were obtained from *G′(ω) = K′* · *ω*^*n*^*′* and *G″(ω) = K″* · *ω*^*n*^*″* by linear regression in log-log space. Conditions were classified as predominantly liquid-like, transitional, or elastic-dominated using tanδ, crossover behavior, the proportion of the frequency window with G′ > G″, and the low-frequency slope of G′.

### 2.3. Critical-gel and thermorheological analyses

The Winter–Chambon criterion was used as a diagnostic rather than as an assumption of ideal gelation. For each sweep, Δn = |n′ − n″|, the coefficient of variation of tanδ, and the deviation between measured tanδ and tan(nmeanπ/2) were calculated. Candidate critical-gel-like conditions were identified by convergence of n′ and n″ together with low frequency dependence of tanδ and agreement between measured and Winter–Chambon-predicted tanδ.

For visualization of the state maps, operational thresholds were defined as liquid-like (tanδ > 1.25), transitional (0.8 ≤ tanδ ≤ 1.25), and elastic-dominated or gel-like (tanδ < 0.8). These thresholds were used as comparative classification boundaries and were interpreted together with crossover behavior and frequency scaling rather than as universal gel-point criteria.

Concentration scaling of G′ and |η*| was evaluated using Y = KCα at representative frequencies and temperatures. Because only three concentrations were available, α was interpreted as an apparent descriptor within the experimental window. Horizontal time-temperature superposition was assessed by shifting G′ and G″ in log-frequency space relative to a central reference temperature. Compatibility required satisfactory overlap, monotonic shift factors, and an approximately linear relationship between log(aT) and 1/T.

Horizontal shift factors were obtained by minimizing the mean squared residual between each shifted G′/G″ sweep and the reference-temperature sweep in log-frequency space. TTS compatibility was evaluated from curve overlap, monotonicity of log(aT), and linearity of log(aT) versus 1/T; conditions with systematic curve-shape changes were classified as thermorheologically complex (Williams et al., 1955).

The dataset architecture and complete analytical workflow are summarized in Figure S3.

### 2.4. Molecular dynamics simulations

Eight GlcNAc10 oligomers were placed in a cubic simulation box as a minimal multichain representation of the highly acetylated fungal EPS. The equilibrated simulation box corresponded to a polymer concentration of approximately 15.8 mg mL^−1^, thereby reproducing the experimental formulation (15 mg mL^−1^) within the uncertainty associated with box equilibration. This model was designed to investigate local ion coordination, short-range chain association, and transient interchain interactions under experimentally relevant polymer concentrations rather than to reproduce the full contour length of the native polymer. Simulations were performed under four conditions (NaCl at 298 and 333 K and LiCl at 298 and 333 K) using the same nominal salt concentrations as the rheological experiments. Systems were built in Schrödinger Materials Science/Maestro, solvated with explicit TIP4P water, and parameterized using the OPLS4 force field (Lu et al., 2021).

After the standard Desmond relaxation protocol, unrestrained NPT production simulations were run for 100 ns at 1.01325 bar. Temperature was controlled using the Nosé-Hoover chain thermostat, pressure using the Martyna-Tobias-Klein barostat, and long-range electrostatics using particle-mesh Ewald. Coordinates were saved every 100 ps, yielding 1001 frames. Three independently initialized trajectories were analyzed per condition, and trajectory-level estimates were summarized as mean ± SD (n = 3 independent simulations) (Bowers et al., 2006).

Custom Python scripts were used to quantify aggregation, compactness, solvent exposure, ion coordination, and interchain bridging. Two chains were assigned to the same cluster when at least three residue-level contacts occurred within 4.5 Å. Radius of gyration (Rg) and polymer solvent-accessible surface area (SASA) described global organization. Ion-polymer radial distribution functions g(r) were calculated between Na^+^ or Li^+^ and polymer oxygen atoms. The first RDF minimum defined the first coordination shell: 3.25 Å for Na+ at both temperatures, 2.85 Å for Li^+^ at 298 K, and 3.05 Å for Li^+^ at 333 K. Continuous coordination events were reconstructed at approximately 1-ns sampling resolution. A cation-mediated interchain bridge was defined when one ion simultaneously coordinated oxygen atoms from at least two distinct chains.

Trajectories were sampled at approx. 1-ns intervals for contact, coordination, and bridge analyses. Consequently, the shortest resolvable continuous event was approx. 1 ns, and reported lifetimes should be interpreted at this temporal resolution.

### 2.5. Statistical analysis

Experimental values at matched conditions were expressed as mean ± SD from three replicate measurements. For MD descriptors, each independent 100-ns trajectory was treated as one computational replicate; frames within a trajectory were not treated as independent observations. Emphasis was placed on effect magnitude, replicate dispersion, and mechanistic consistency rather than pseudo-replication across temporally correlated frames. Analyses and figures were generated using Python and GraphPad Prism.

## 3. Results and discussion

### 3.1. Concentration defines distinct viscoelastic regimes

The oscillatory profiles established concentration as the principal control of the rheological state (Figure 1). At 5 mg mL^−1^, both ionic media displayed low moduli and strong frequency dependence, consistent with weakly associated solutions. At 10 mg mL^−1^, elastic contributions became measurable but remained salt dependent. At 15 mg mL^−1^, G′ dominated broader portions of the frequency window, indicating a more solid-like response and a persistent physical network.

**Figure 1.**
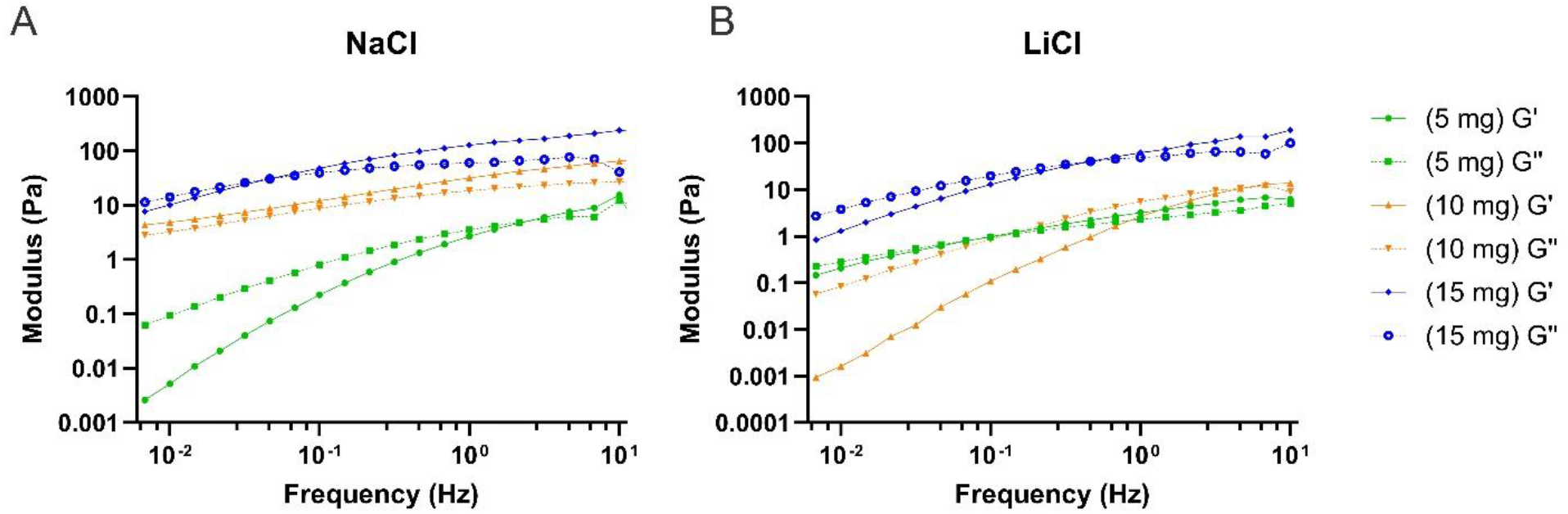
Frequency-dependent viscoelastic behavior of fungal chitin-like EPS solutions. Representative oscillatory frequency sweeps recorded at approximately 15 °C for EPS solutions prepared in (A) NaCl and (B) LiCl at 5, 10, and 15 mg mL^−1^. Solid and dashed curves represent replicate-averaged G′ and G″ values, respectively (n = 3). Data are shown over the reliable analytical frequency range (0.01–10 Hz). Representative numerical values and associated standard deviations are provided in Table S2.

The two salts nevertheless generated different rheological responses. At 10 mg mL^−1^, LiCl produced high absolute moduli but a predominantly dissipative response, as G″ increased in concomitance with G′. In contrast, NaCl produced lower absolute moduli under some conditions but lower tanδ values and more frequent G′ dominance over a broader frequency range. These results highlight why modulus magnitude alone is insufficient to characterize gel-like behavior: the viscoelastic response depends on the relative contributions of stored and dissipated energy, rather than solely on the magnitude of G′.

Apparent concentration-scaling exponents supported this interpretation. At approximately 25 °C and 1 Hz, NaCl showed steeper and more coherent increases in |η*| and G′ with concentration than LiCl (Figure S1). Because the exponents were based on three concentration levels, they are best interpreted as formulation-range descriptors rather than universal scaling laws. The apparent concentration-scaling and TTS descriptors are summarized in Table S3.

### 3.2. Temperature and salt identity shift the network response

Temperature did not act as a simple thinning variable (Figure 2). Changes in G′, G″, and |η*| depended on concentration and salt, indicating thermal reorganization of the relaxation spectrum. The contrast was clearest in concentrated systems. NaCl showed a biphasic or reinforcing response in which G′ recovered or increased after an intermediate-temperature minimum, whereas LiCl generally softened with increasing temperature.

**Figure 2.**
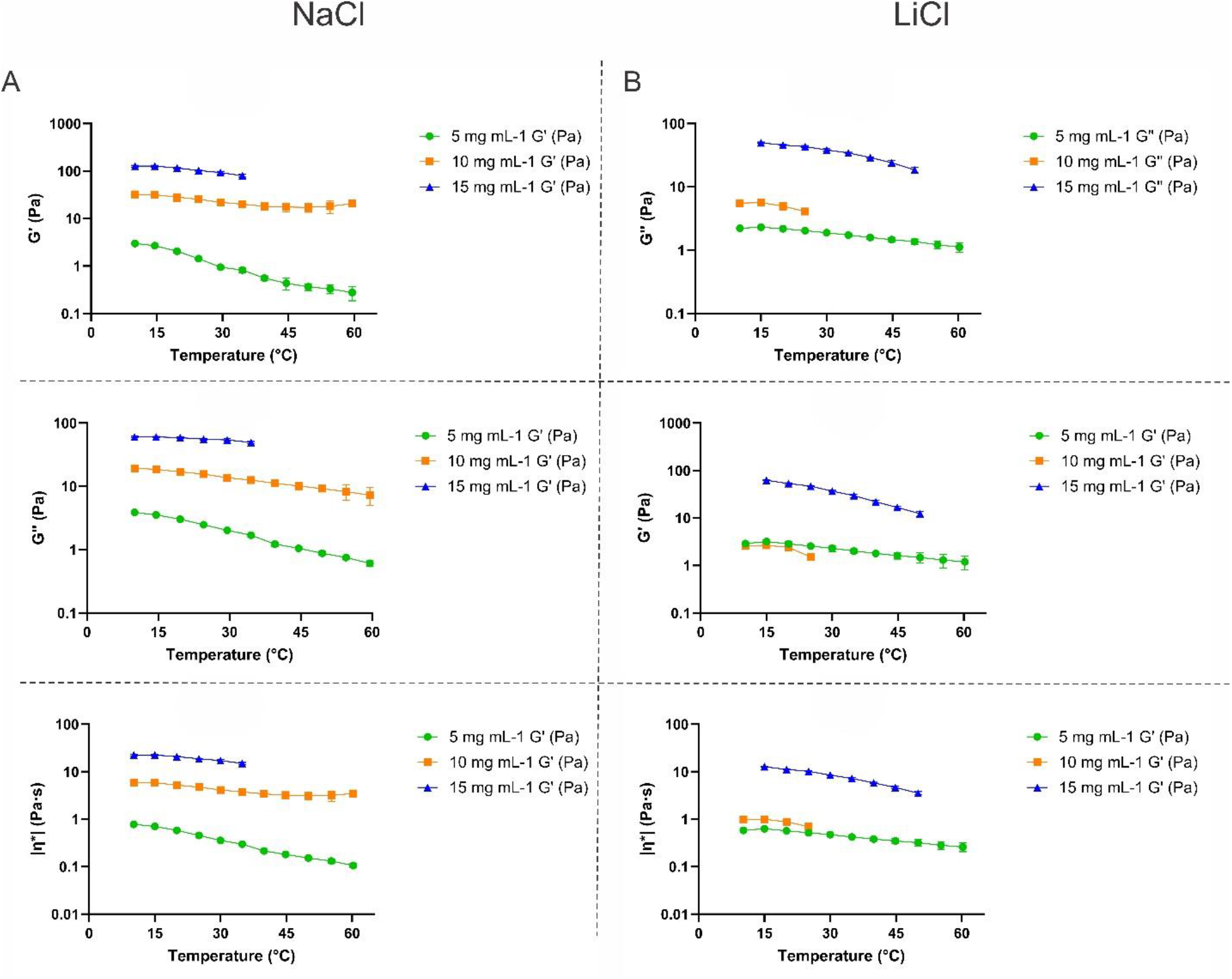
Temperature-dependent modulation of the viscoelastic response. Temperature dependence of G′ (A,B), G″ (C,D), and |η*| (E,F), evaluated at approximately 1 Hz for EPS solutions prepared in NaCl and LiCl at 5, 10, and 15 mg mL^−1^. Values represent mean ± SD from three replicate measurements at matched conditions. Temperature ranges reflect the available experimental measurements for each formulation; no interpolation was applied to conditions for which measurements were unavailable.

Apparent Arrhenius analysis provided comparative measures of temperature sensitivity but was not universally applicable. Several formulations showed satisfactory linearity, whereas the 10 mg mL^−1^ LiCl system did not follow a simple Arrhenius relationship. Similarly, time-temperature superposition was strongest for the 15 mg mL^−1^ NaCl system and only partial or conditionally valid elsewhere (Figure S2). Linear regression of ln(aT) against reciprocal absolute temperature further supported Arrhenius-type shift-factor behavior for NaCl at 15 mg mL^−1^ (R^2^ = 0.971, p = 0.0021), LiCl at 15 mg mL^−1^ (R^2^ = 0.967, p < 0.0001), and NaCl at 10 mg mL^−1^ (R^2^ = 0.891, p = 0.0158). Although LiCl at 10 mg mL^−1^ showed a high apparent R^2^ (0.957), the slope was not significantly different from zero (p = 0.133), reflecting the limited number of available temperature points. These results indicate that Arrhenius-like temperature dependence of the horizontal shift factors was formulation-dependent rather than universal.

Failure of global superposition is mechanistically meaningful because it indicates that heating changed network topology rather than merely shifting relaxation times.

### 3.3. tanδ mapping and critical-gel diagnostics identify salt-specific transition pathways

State maps based on tanδ summarized the combined influence of concentration and temperature (Figure 3). NaCl displayed a comparatively direct progression from liquid-like behavior at low concentration toward elastic-dominated behavior at 10 and 15 mg mL^−1^. LiCl followed a non-monotonic pathway, including a highly dissipative intermediate regime at 10 mg mL^−1^ before approaching lower tanδ at the highest concentration.

**Figure 3.**
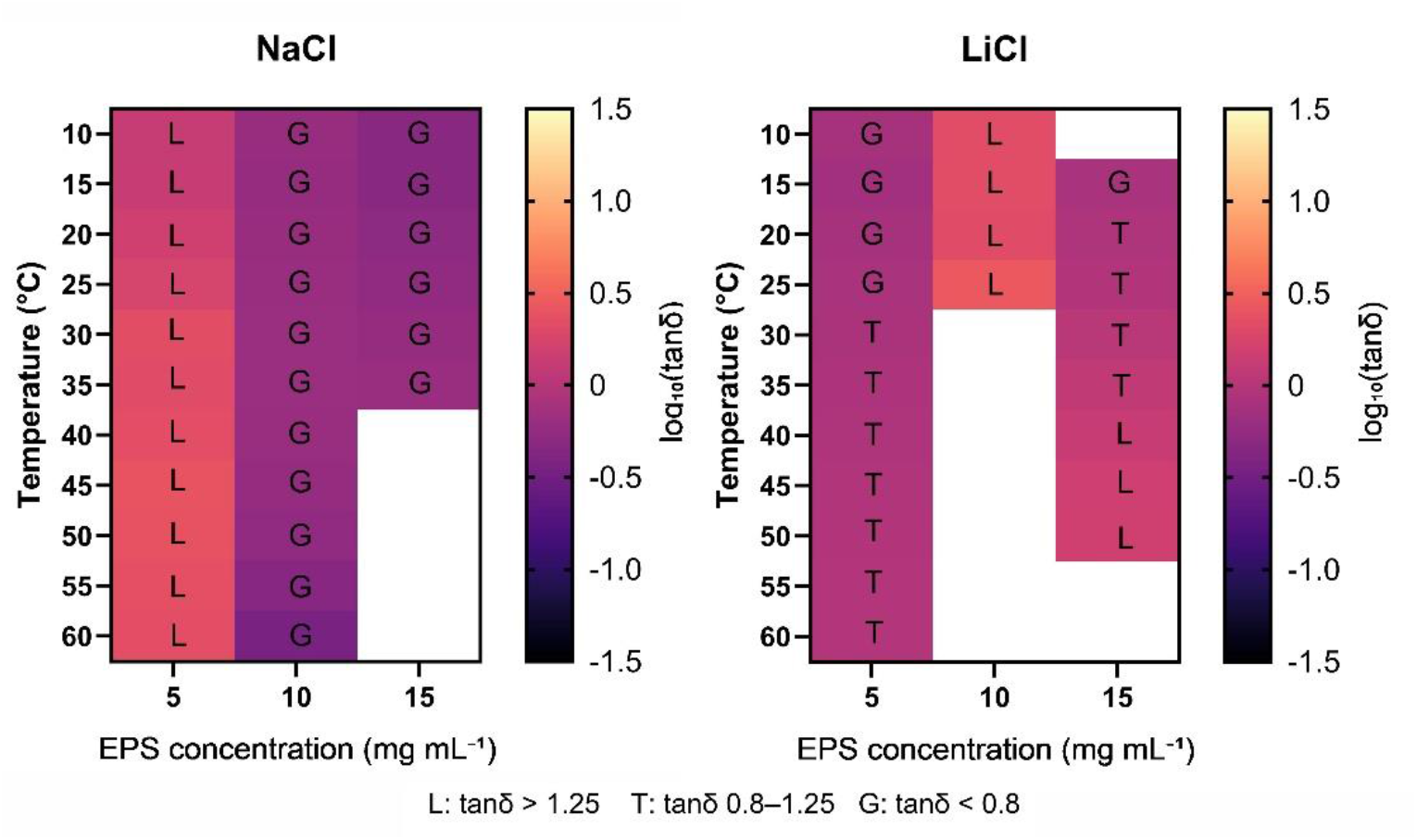
tanδ-based rheological state maps. Concentration-temperature maps of log10(tanδ) at approximately 1 Hz for NaCl and LiCl solutions. L, liquid-like (tanδ > 1.25); T, transitional (0.8 ≤ tanδ ≤ 1.25); G, elastic-dominated or gel-like (tanδ < 0.8). White cells indicate unavailable or excluded measurements.

Winter–Chambon diagnostics identified regions that approached, but did not perfectly satisfy, ideal critical-gel behavior (Figure 4). The most informative conditions combined convergence of n′ and n″, low frequency dependence of tanδ, and agreement between measured and theoretically predicted tanδ. We therefore use the terms gel-like and critical-gel-like cautiously. The data support a dynamic physical network and a percolation-like increase in connectivity, not a permanently crosslinked equilibrium gel.

**Figure 4.**
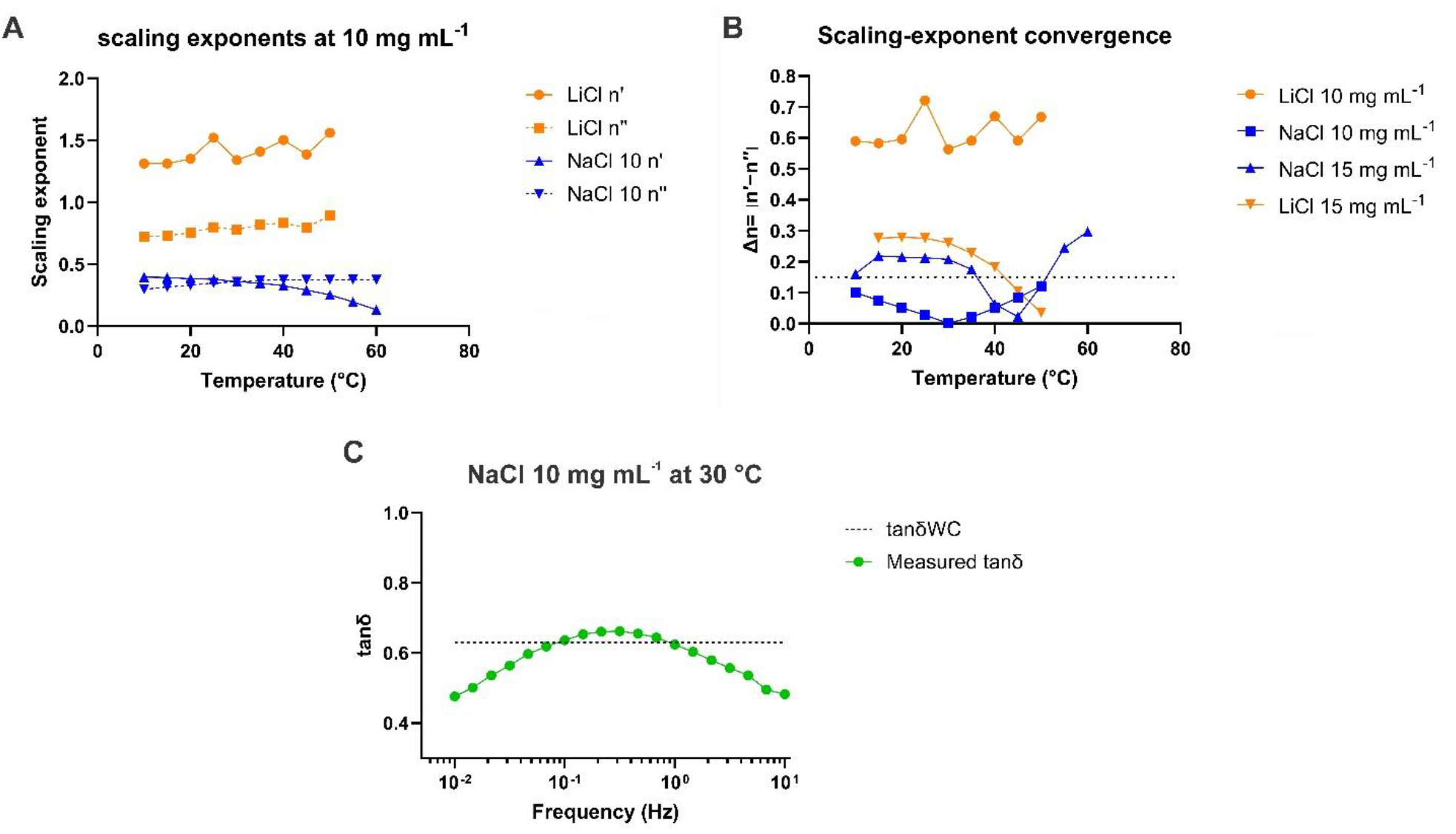
Winter–Chambon diagnostic analysis. (A) Representative scaling exponents n′ and n″. (B) Convergence of the scaling exponents, expressed as Δn = |n′ − n″|. (C) Comparison of measured and Winter–Chambon-predicted tanδ for a candidate critical-gel-like condition. Smaller Δn values and weaker frequency dependence of tanδ indicate closer agreement with ideal critical-gel behavior.

### 3.4. Molecular configurations reveal opposing assembly responses

The molecular snapshots provide a direct visual comparison between the initially dispersed oligomers and the final configurations (Figure 5). The representative configurations visually suggest more frequent chain–chain association in NaCl, consistent with the replicate-averaged descriptors in Figure 6.

**Figure 5.**
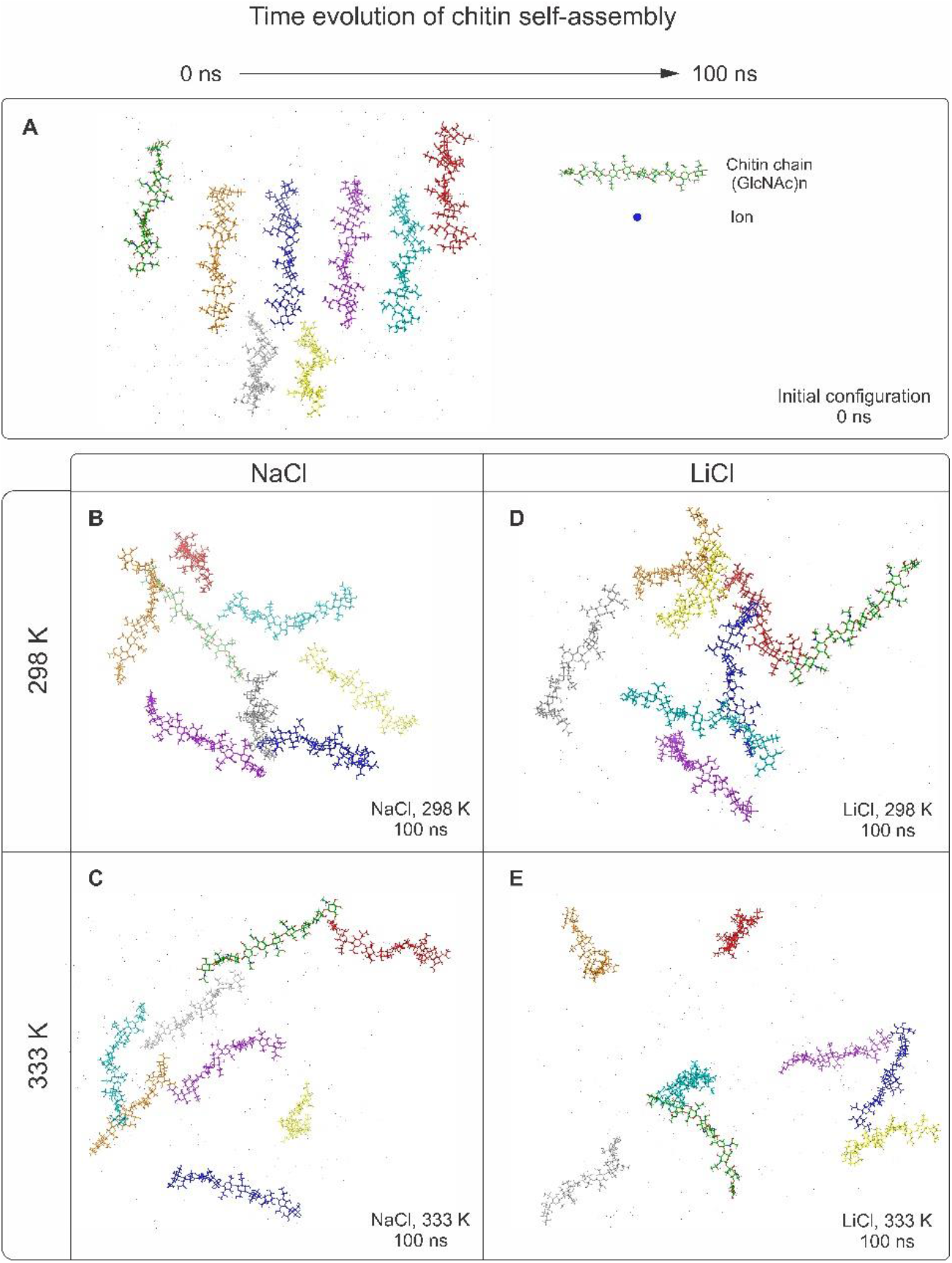
Initial and final molecular organization of chitin chain GlcNAc10 oligomers under different ionic and thermal conditions. (A) Representative initial configuration at 0 ns. Final configurations after 100 ns are shown for (B) NaCl at 298 K, (C) NaCl at 333 K, (D) LiCl at 298 K, and (E) LiCl at 333 K. Each oligomer is displayed in a distinct color and retains the same color across panels to facilitate visual tracking. The snapshots are shown for qualitative visualization; quantitative comparisons are presented in Figures 6–9.

**Figure 6.**
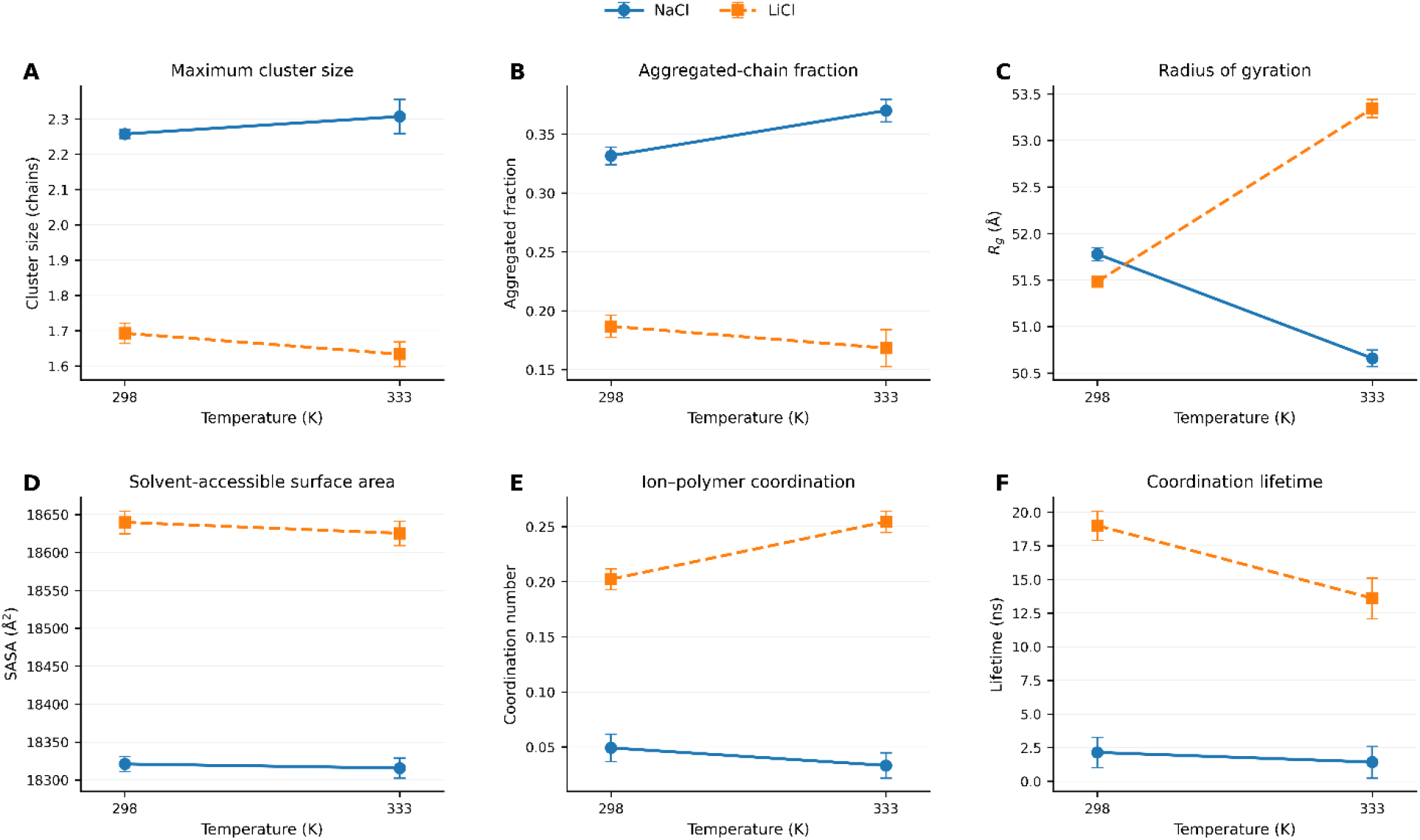
Temperature- and salt-dependent molecular organization of the fungal chitin-like EPS. (A) Mean maximum cluster size. (B) Aggregated-chain fraction. (C) Radius of gyration. (D) Polymer SASA. (E) Mean ion-polymer coordination number. (F) Mean coordination lifetime. Points represent the mean of three independent 100-ns simulations; error bars indicate ±SD.

Heating produced opposing responses in the two salts. In NaCl, the mean maximum cluster size and aggregated-chain fraction increased with the temperature, while Rg decreased from 51.779 ± 0.069 to 50.660 ± 0.090 Å (Figure 6A-C). In LiCl, cluster size and aggregated fraction decreased, while Rg increased from 51.483 ± 0.011 to 53.345 ± 0.098 Å. SASA changed little, indicating that the principal response involved chain packing and connectivity rather than wholesale collapse or dehydration. Trajectory-level global assembly descriptors are summarized in Table S4.

### 3.5. Local ion coordination is decoupled from collective aggregation

The RDF profiles revealed much stronger localization of Li^+^ around polymer oxygen atoms than of Na^+^ (Figure 7). The Na^+^-O peak occurred at 2.55 Å at 298 K and 2.45 Å at 333 K, with first minima at 3.25 Å and coordination numbers of 0.050 and 0.033. The Li^+^-O peak remained at 2.05 Å, but its intensity increased at 333 K; the first minimum shifted from 2.85 to 3.05 Å and the coordination number increased from 0.202 to 0.255.

**Figure 7.**
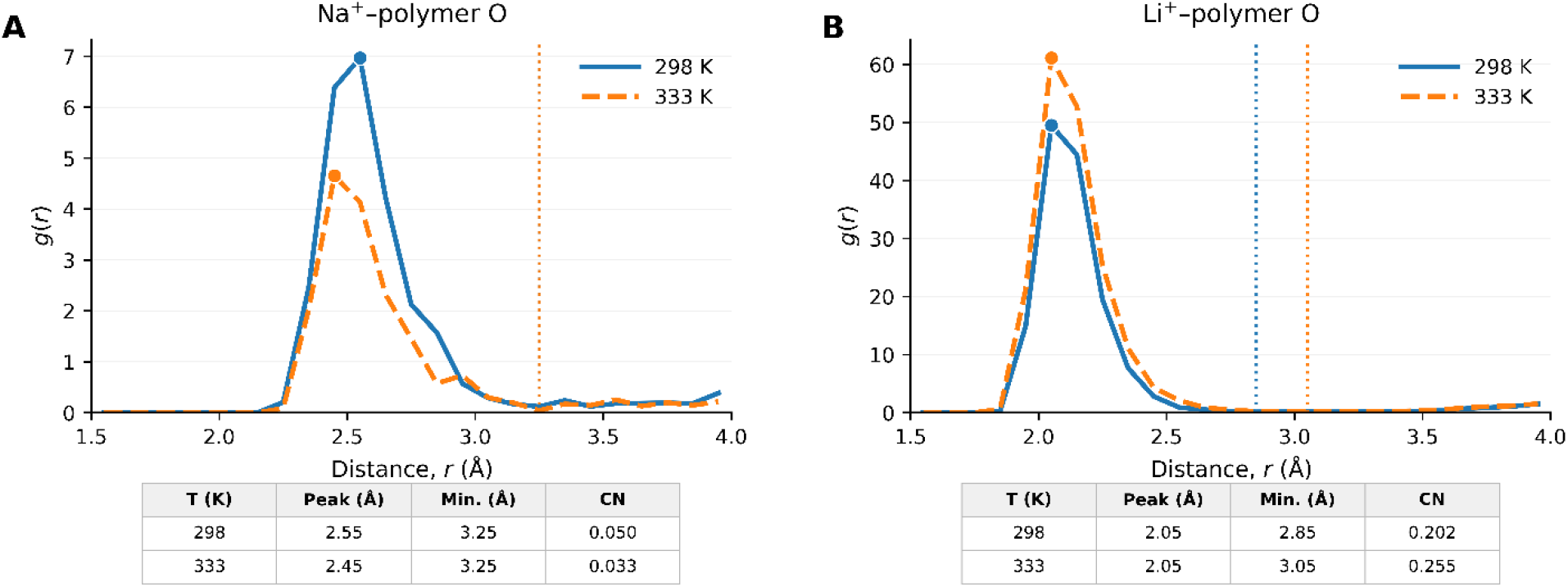
First-shell ion-polymer radial distribution functions. (A) Na^+^-polymer oxygen RDFs at 298 and 333 K. (B) Li^+^-polymer oxygen RDFs at 298 and 333 K. Symbols indicate the principal maxima and dotted vertical lines the first minima used as coordination cutoffs. Peak positions, minima, and integrated coordination numbers are reported below each panel. RDFs were calculated independently for each of the three 100-ns trajectories and subsequently averaged for each condition (n = 3).

Coordination distributions confirmed that Li^+^ sampled coordinated states more frequently and maintained them for longer times (Figure 8). Mean Li^+^ coordination lifetimes decreased from 19.01 to 13.61 ns with heating but remained far longer than the corresponding Na^+^ lifetimes of 2.14 and 1.43 ns. Heating therefore accelerated exchange in both salts, while increasing the instantaneous population of Li+-coordinated states. Numerical RDF and first-shell coordination descriptors are provided in Table S5.

**Figure 8.**
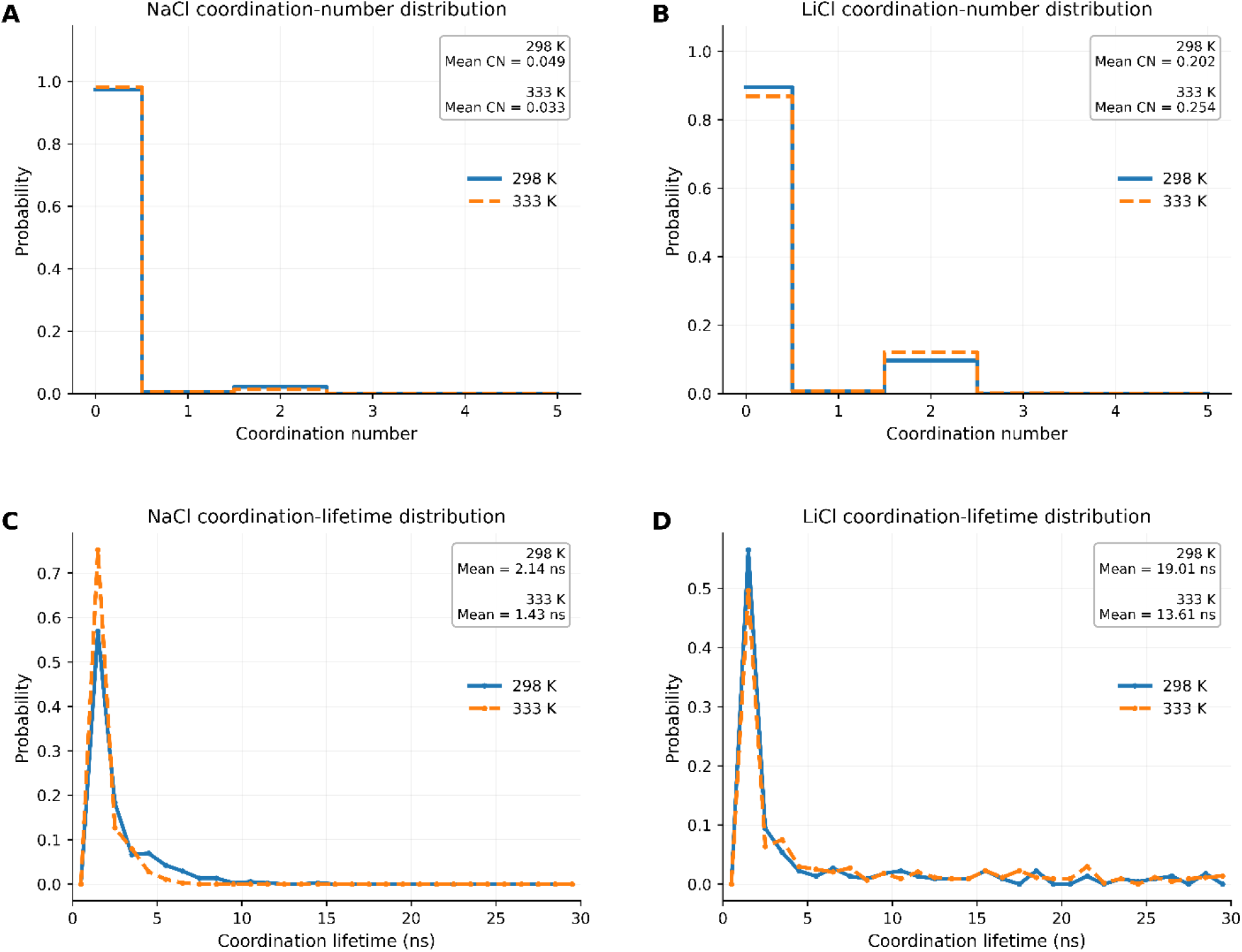
Distributions of ion-polymer coordination number and coordination lifetime. Coordination-number probability distributions for (A) NaCl and (B) LiCl. Coordination-lifetime probability distributions for (C) NaCl and (D) LiCl. Solid and dashed curves represent 298 and 333 K, respectively. Coordination-number and coordination-lifetime distributions were calculated independently for each trajectory and averaged across three independent 100-ns simulations for each condition (n = 3).

The central mechanistic result is that stronger local coordination did not produce greater global aggregation. LiCl generated the sharpest first-shell structure and longest residence times, yet its oligomers formed smaller and more expanded assemblies. This is consistent with the compact hydration environment and high charge density of Li^+^, which can stabilize localized polymer-ion-water configurations (Mason et al., 2015). Na^+^ interacted more weakly and dynamically, potentially allowing chain packing and polymer-polymer contacts to reorganize more cooperatively.

### 3.6. Interchain ion bridges are transient and do not alone define network strength

Direct cation-mediated bridges between two or more chains were sparse and short-lived (Figure 9). NaCl produced three events at 298 K and five at 333 K, with average lifetimes of approximately 1 ns. No Li^+^-mediated bridges were detected at 298 K under the applied criteria, whereas nine were detected at 333 K with an average lifetime of 1.11 ns. These events were intermittent rather than continuously occupied cross-links.

**Figure 9.**
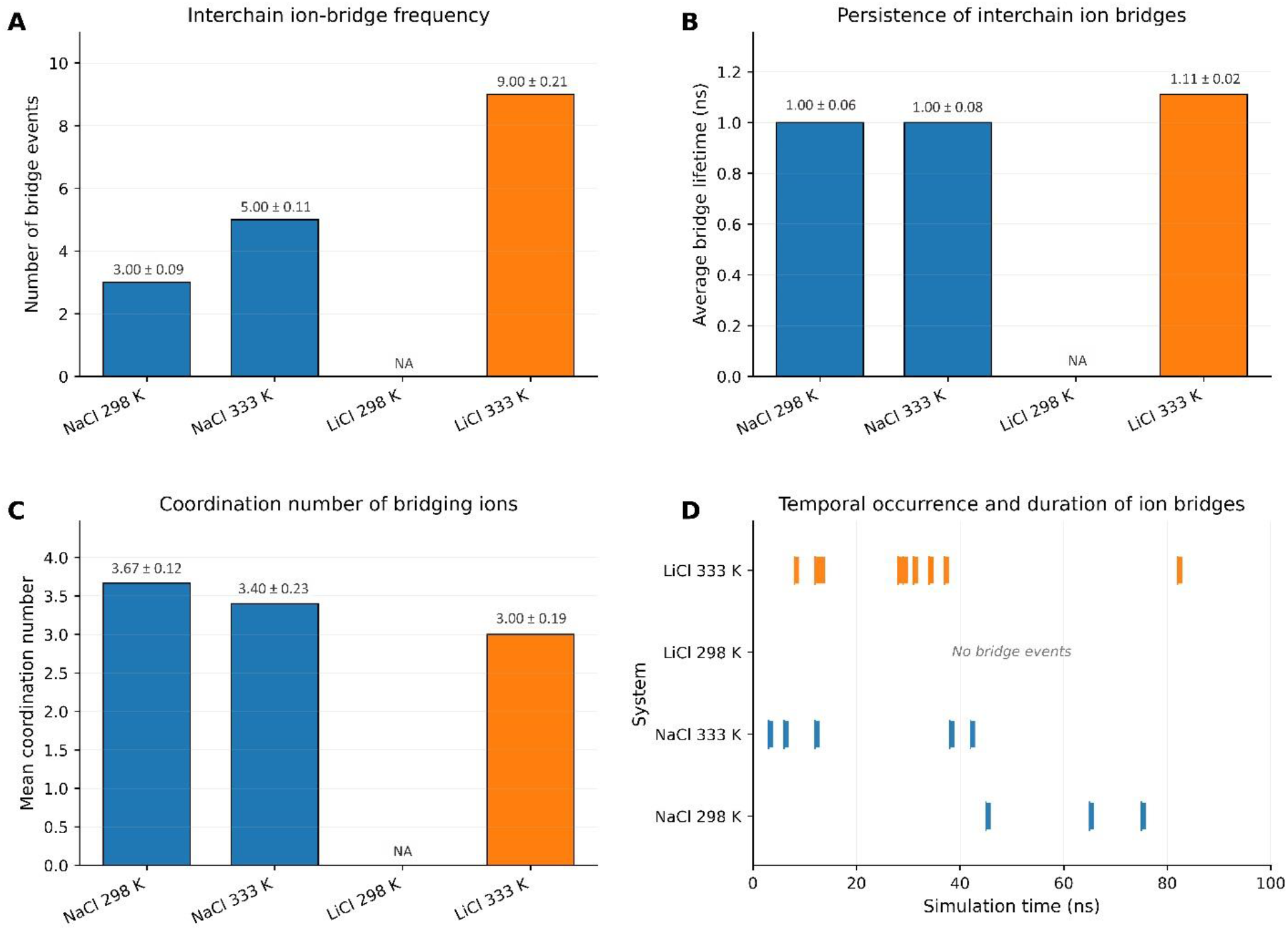
Transient ion-mediated interchain bridging. (A) Number of bridge events, (B) bridge lifetime, (C) coordination number of bridging ions, and (D) representative temporal occurrence and duration of bridge events during 100-ns MD simulations. Values in A–C are mean ± SD from three independent trajectories (n = 3). No Li^+^-mediated interchain bridges were detected at 298 K; therefore, lifetime and coordination number are reported as N/A.

The greater number of Li^+^ bridge events at 333 K did not correspond to a more compact global assembly. Bridge count should therefore not be equated with mechanical percolation. The events instead reflect rapid sampling of multichain geometries within a locally coordinated but globally expanded system. In NaCl, fewer direct cation bridges were compatible with stronger collective aggregation, suggesting that ionic screening and rapid exchange facilitate polymer proximity while polymer-polymer hydrogen-bond-like contacts and packing interactions provide a larger share of network connectivity. Interchain bridge counts and lifetime descriptors are summarized in Table S6.

Together, the experimental and computational findings support a hierarchical mechanism. Increasing concentration raises chain overlap and makes interchain association mechanically relevant. Salt identity then determines how local coordination couples to collective packing. NaCl behaves primarily as a dynamic modulator of chain association and favors compact, connected assemblies; LiCl acts as a stronger local coordination center but can stabilize more solvated, spatially expanded chain-ion configurations. The observed gel-like response is consequently a reversible physical network governed by the population and exchange of multiple weak interactions rather than permanent ionic cross-links.

Although LiCl at 333 K produced a higher absolute number of bridge events, normalization by cation abundance yielded a rate comparable to that of NaCl at the same temperature (3.23 and 3.25 events per 100 cations per 100 ns, respectively). Thus, the raw bridge count mainly reflects the larger number of Li^+^ ions in the simulated system and should not be interpreted as stronger network formation. In the native high-molecular-weight EPS, however, multiple transient bridges distributed along much longer chains could collectively contribute to network connectivity, an effect that cannot be captured by the present GlcNAc10 model.

### 3.7. Implications and limitations

The combined analysis identifies ionic formulation as a formulation variable that controls the route by which local molecular interactions are converted into macroscopic viscoelasticity. The result is relevant to processing because a formulation may display high modulus while remaining highly dissipative, or comparatively modest local ion binding while developing a coherent elastic network.

Salt-specific differences between local ionic association and macroscopic gel reinforcement have also been observed in other polysaccharide systems, although the molecular origin and charge state of the present highly acetylated chitin-like EPS are distinct (Yang et al., 2024).

Several limitations should be considered. The experimental analysis reuses an existing rheological dataset and depends on the completeness of the original replicate structure. Apparent scaling exponents are based on three concentrations. Although the simulations reproduced the experimental polymer mass concentration (∼15 mg mL^−1^), they employed a minimal multichain model composed of eight GlcNAc10 oligomers rather than the native fungal EPS (Mw ≈ 4.9 × 10^5^ g mol^−1^). Consequently, the simulations were intended to resolve local ion coordination, short-range chain association, and transient interchain bridging, rather than long-chain conformational, entanglement, or percolation behavior. This complementary multiscale strategy allows molecular interactions to be interpreted in the context of experimentally observed viscoelastic behavior without claiming an atomistic reconstruction of the macroscopic gel network.

The reconstituted EPS solutions were under near-neutral conditions and no deliberate pH adjustment was performed. Given the high degree of acetylation (>90%), only a minor fraction of glucosamine units contains potentially protonatable amino groups, and their contribution to the overall chain charge is therefore expected to be limited. Nevertheless, a minor contribution of residual protonation to chain conformation, hydration, and intermolecular interactions cannot be completely excluded.

In addition, NaCl and LiCl were evaluated at different molar concentrations (0.154 and 0.28 mol L^-1^, respectively). Consequently, the observed differences reflect the complete ionic formulations and cannot be attributed exclusively to cation identity or hydration chemistry.

## 4. Conclusions

Fungal chitin-like EPS solutions exhibited a salt-dependent thermorheological response governed by polymer concentration, temperature, and oscillation timescale. Increasing EPS concentration shifted the systems from predominantly liquid-like behavior toward transitional and elastic-dominated regimes, with NaCl promoting a more direct progression toward network connectivity than LiCl. Molecular dynamics simulations revealed that this difference did not arise simply from the strength of direct cation-polymer binding. NaCl favored larger and more compact polymer assemblies despite relatively weak and short-lived Na^+^-oxygen coordination, whereas LiCl produced stronger and more persistent local Li^+^-oxygen interactions while maintaining smaller, more expanded assemblies. Ion-mediated interchain bridges were sparse and transient, supporting a dynamic physical-network mechanism rather than permanent ionic crosslinking. Overall, the NaCl and LiCl formulations differed in how local ion coordination was coupled to collective polymer association, showing that stronger local ion binding does not necessarily translate into greater macroscopic network reinforcement. This multiscale relationship provides molecular insight into how ionic conditions can be used to tune the viscoelastic response of fermentation-derived chitin-like EPS systems.

## CRediT authorship contribution statement

Barazorda Ccahuana: Data curation, Formal analysis, Writing - original draft, Writing - review and editing. Rilton Alves de Freitas: Investigation, Methodology, Data curation. Júlio Cesar de Carvalho: Conceptualization, Supervision, Resources. Miguel Daniel Noseda: Methodology, Validation, Resources. Carlos Ricardo Soccol: Conceptualization, Resources, Supervision. Luis Daniel Goyzueta-Mamani: Conceptualization, Formal analysis, Software, Visualization, Writing - original draft, Writing - review and editing, Project administration.

## Acknowledgments

M.D.N., R.A.F., C.R.S., J.C.C. are Research Members of the National Research Council of Brazil (CNPq).

## Data availability

The processed rheological descriptors, trajectory-level molecular dynamics summaries, and scripts supporting the findings of this study will be made available in a public repository upon acceptance. The original EPS characterization and rheological dataset are described in (Goyzueta M. et al., 2020).

## References

Bowers, K. J., Chow, E., Xu, H., Dror, R. O., Eastwood, M. P., Gregersen, B. A., Klepeis, J. L., Kolossvary, I., Moraes, M. A., Sacerdoti, F. D., Salmon, J. K., Shan, Y., & Shaw, D. E. (2006). Scalable algorithms for molecular dynamics simulations on commodity clusters. Proceedings of the 2006 ACM/IEEE Conference on Supercomputing, 84–es. 10.1145/1188455.1188544

Chambon, F., & Winter, H. H. (1987). Linear viscoelasticity at the gel point of a crosslinking PDMS with imbalanced stoichiometry. Journal of Rheology, 31(8), 683–697.

Chen, Y., Zhu, H., Feng, Y., Wang, Y., Wang, X., Zhang, W., & Luo, Y. (2026). Rheological analysis in food processing: factors, applications, and future outlooks with machine learning integration. RSC Advances, 16(6), 5040–5063.

Goyzueta M. L. D., Noseda, M. D., Bonatto, S. J. R., de Freitas, R. A., de Carvalho, J. C., & Soccol, C. R. (2020). Production, characterization, and biological activity of a chitin-like EPS produced by Mortierella alpina under submerged fermentation. Carbohydrate Polymers, 116716. 10.1016/j.carbpol.2020.116716

Kaur, S., & Dhillon, G. S. (2015). Recent trends in biological extraction of chitin from marine shell wastes: a review. Critical Reviews in Biotechnology, 35(1), 44–61.

Lu, C., Wu, C., Ghoreishi, D., Chen, W., Wang, L., Damm, W., Ross, G. A., Dahlgren, M. K., Russell, E., & Von Bargen, C. D. (2021). OPLS4: Improving force field accuracy on challenging regimes of chemical space. Journal of Chemical Theory and Computation, 17(7), 4291–4300.

Mähler, J., & Persson, I. (2011). A Study of the Hydration of the Alkali Metal Ions in Aqueous Solution. Inorganic Chemistry, 51(1), 425–438. 10.1021/ic2018693

Mason, P. E., Ansell, S., Neilson, G. W., & Rempe, S. B. (2015). Neutron scattering studies of the hydration structure of Li+. The Journal of Physical Chemistry. B, 119(5), 2003–2009. 10.1021/jp511508n

McGregor, N. G. S., Penttilä, P., Pitkänen, L., Mohammadi, P., Vuorte, M., Igarashi, K., & Arola, S. (2025). Self-assembly of mixed-linkage glucan hydrogels formed following EG16 digestion. Carbohydrate Polymers, 347, 122703. 10.1016/j.carbpol.2024.122703

Narkevicius, A., Steiner, L. M., Parker, R. M., Ogawa, Y., Frka-Petesic, B., & Vignolini, S. (2019). Controlling the Self-Assembly Behavior of Aqueous Chitin Nanocrystal Suspensions. Biomacromolecules, 20(7), 2830–2838. 10.1021/acs.biomac.9b00589

Picout, D. R., & Ross-Murphy, S. B. (2003). Rheology of Biopolymer Solutions and Gels. The Scientific World Journal, 3(1), 524215. 10.1100/tsw.2003.15

Pieklarz, K., Jenczyk, J., Modrzejewska, Z., Owczarz, P., & Jurga, S. (2022). An Investigation of the Sol-Gel Transition of Chitosan Lactate and Chitosan Chloride Solutions via Rheological and NMR Studies. In Gels (Vol. 8, Issue 10, p. 670). 10.3390/gels8100670

Pillai, C. K. S., Paul, W., & Sharma, C. P. (2009). Chitin and chitosan polymers: Chemistry, solubility and fiber formation. Progress in Polymer Science (Oxford), 34(7), 641–678. 10.1016/j.progpolymsci.2009.04.001

Rinaudo, M. (2006). Chitin and chitosan: Properties and applications. Progress in Polymer Science (Oxford), 31(7), 603–632. 10.1016/j.progpolymsci.2006.06.001

Rinaudo, M., Milas, M., & Dung, P. Le. (1993). Characterization of chitosan. Influence of ionic strength and degree of acetylation on chain expansion. International Journal of Biological Macromolecules, 15(5), 281–285. 10.1016/0141-8130(93)90027-J

Romany, A., Payne, G. F., & Shen, J. (2023). Mechanism of the Temperature-Dependent Self-Assembly and Polymorphism of Chitin. Chemistry of Materials, 35(16), 6472–6481. 10.1021/acs.chemmater.3c01313

Steinfeld, L., Vafaei, A., Rösner, J., & Merzendorfer, H. (2019). Chitin prevalence and function in bacteria, fungi and protists. Targeting Chitin-Containing Organisms, 19–59.

Stojkov, G., Niyazov, Z., Picchioni, F., & Bose, R. K. (2021). Relationship between Structure and Rheology of Hydrogels for Various Applications. Gels (Basel, Switzerland), 7(4). 10.3390/gels7040255

Synowiecki, J., & Al-Khateeb, N. A. (2003). Production, properties, and some new applications of chitin and its derivatives. Critical Reviews in Food Science and Nutrition, 43(2), 145–171. 10.1080/10408690390826473

Teng, W. L., Khor, E., Tan, T. K., Lim, L. Y., & Tan, S. C. (2001). Concurrent production of chitin from shrimp shells and fungi. Carbohydrate Research, 332(3), 305–316.

Williams, M. L., Landel, R. F., & Ferry, J. D. (1955). The temperature dependence of relaxation mechanisms in amorphous polymers and other glass-forming liquids. Journal of the American Chemical Society, 77(14), 3701–3707.

Winter, H. H., & Chambon, F. (1986). Analysis of linear viscoelasticity of a crosslinking polymer at the gel point. Journal of Rheology, 30(2), 367–382.

Winter, H. H., & Mours, M. (1999). Rheology of polymers near liquid-solid transitions. In Neutron spin echo spectroscopy viscoelasticity rheology (pp. 165–234). Springer.

Yang, X., Kimura, M., Zhao, Q., Ryo, K., Descallar, F. B. A., & Matsukawa, S. (2024). Gelation of gellan induced by trivalent cations and coexisting trivalent with monovalent cations studied by rheological and DSC measurements. Carbohydrate Polymers, 345, 122485. 10.1016/j.carbpol.2024.122485

Younes, I., & Rinaudo, M. (2015). Chitin and chitosan preparation from marine sources. Structure, properties and applications. Marine Drugs, 13(3), 1133–1174. 10.3390/md13031133

Zhao, Y., Cao, Y., Yang, Y., & Wu, C. (2003). Rheological Study of the Sol−Gel Transition of Hybrid Gels. Macromolecules, 36(3), 855–859. 10.1021/ma020919y

