## Supplementary material for "Thermorheological mapping and molecular insights into salt-dependent gel-like network formation in *Mortierella alpina* chitin-like exopolysaccharide solutions"

### Contents

Figure S1. Concentration-scaling analysis

Figure S2. Time-temperature superposition diagnostics

Figure S3. Dataset architecture and analytical workflow

Figure S4. Complete crossover and tanδ maps

Table S1. Experimental design and replicate structure

Table S2. Representative rheological descriptors

Table S3. Apparent concentration-scaling and thermorheological parameters

Table S4. Global MD descriptors

Table S5. RDF and ion-coordination descriptors

Table S6. Interchain bridge descriptors

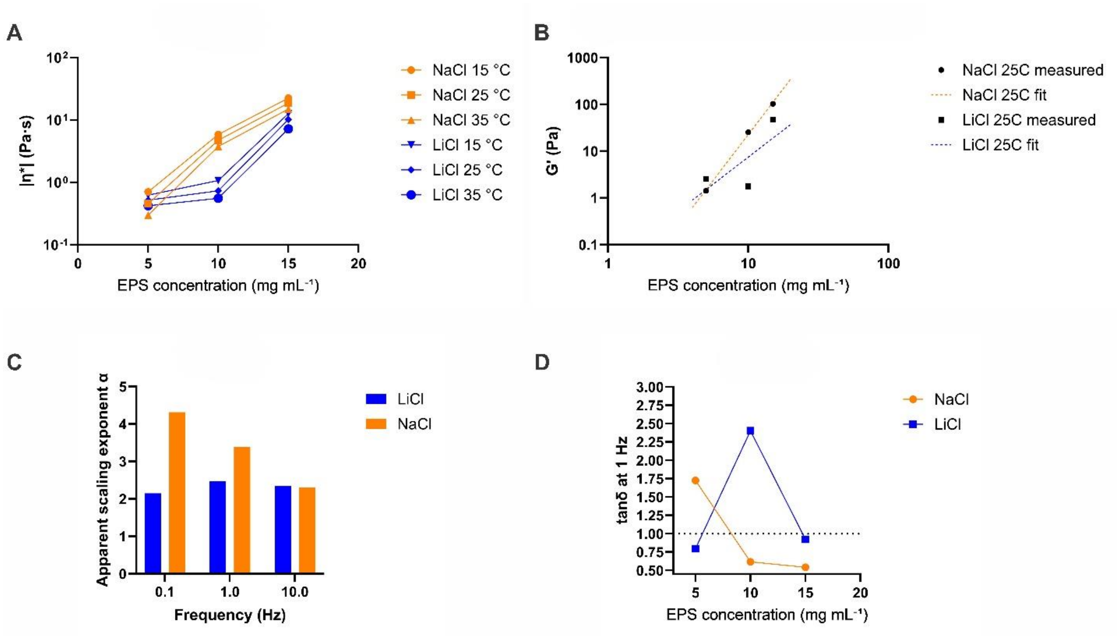


**Figure S1**. Concentration-scaling analysis. (A) Complex viscosity as a function of EPS concentration at 1 Hz. (B) Power-law concentration scaling of G′ at approximately 25 °C. (C) Apparent concentration-scaling exponent of |η*| at 0.1, 1, and 10 Hz. (D) tanδ as a function of concentration at 1 Hz and approximately 25 °C. Exponents are formulation-range descriptors based on three concentration levels.

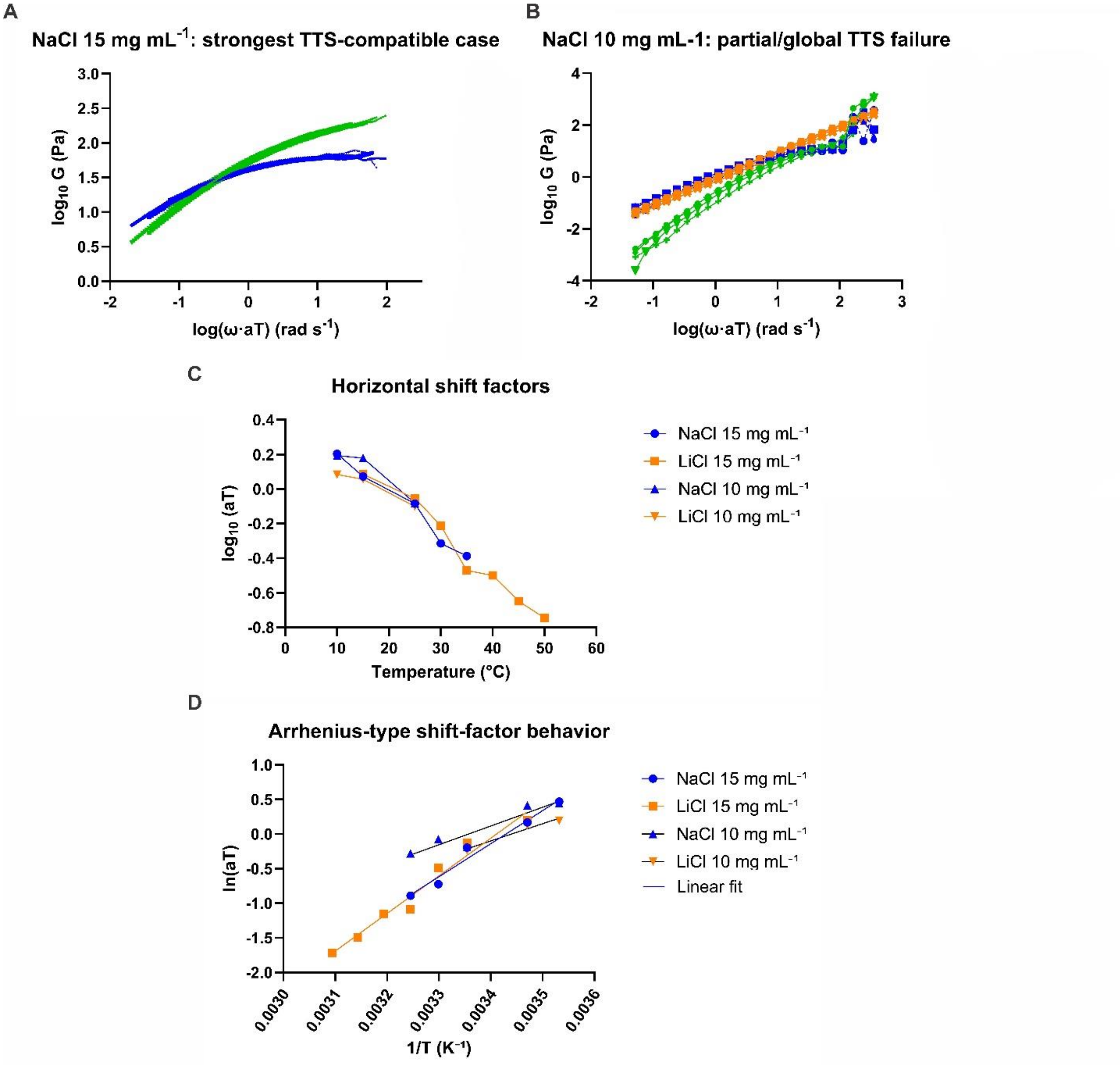


**Figure S2.** Time–temperature superposition (TTS) analysis of EPS solutions. (A) Master curves of G′ and G″ for NaCl 15 mg mL⁻¹, showing the strongest TTS-compatible behavior. (B) Frequency-shifted data for NaCl 10 mg mL⁻¹, illustrating incomplete TTS. (C) Horizontal shift factors, $\log_{10}$(aT), for NaCl and LiCl systems at 10 and 15 mg mL⁻¹. (D) Arrhenius-type representation of the shift factors as ln(aT) versus 1/T; lines represent linear regressions.

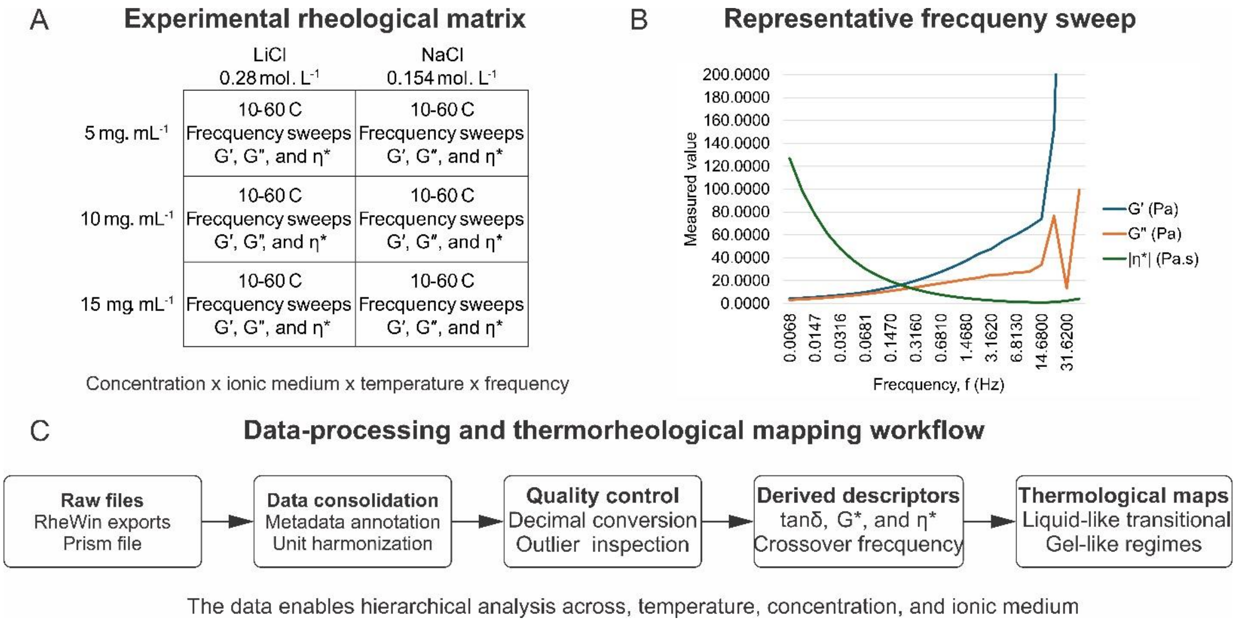


**Figure S3**. Architecture of the rheological dataset and analytical workflow. (A) Experimental matrix. (B) Representative frequency sweep. (C) Data consolidation, quality control, calculation of derived descriptors, and generation of rheological state maps.

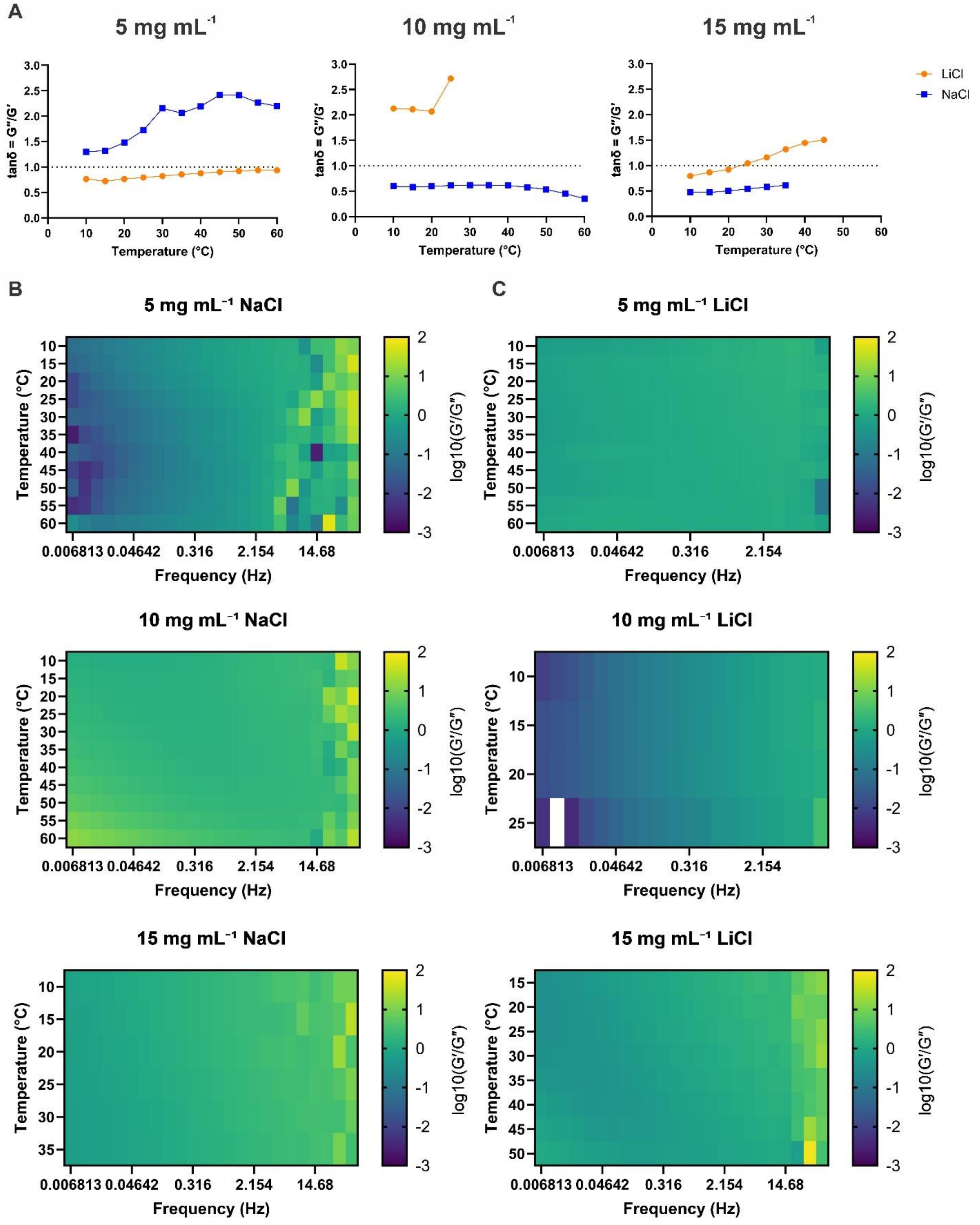


**Figure S4. Complete crossover and modulus-ratio maps.** (A) Representative tanδ profiles as a function of temperature. (B–C) Concentration-, temperature-, and frequency-dependent maps of $\log_{10}(G'/G'')$ for the NaCl and LiCl formulations.

**Supplementary tables**

**Table S1.** Experimental design and replicate structure.

| Condition | Temperature range (°C) | Replicates | Maximum frequency (Hz) | Notes |
|---|---|---|---|---|
| 5 mg mL$^{-1}$ NaCl | 10-60 | 3 | 46.42 | Full retained range |
| 5 mg mL$^{-1}$ LiCl | 10-60 | 3 | 46.42 | Full retained range |
| 10 mg mL$^{-1}$ NaCl | 10-60 | 3 | 46.42 | Full retained range |
| 10 mg mL$^{-1}$ LiCl | 10-60 | 3 | 46.42 | Full retained range |
| 15 mg mL$^{-1}$ NaCl | 10-60 | 3 | 46.42 | Full retained range |
| 15 mg mL$^{-1}$ LiCl | 15-50 | 3 | 46.42 | Data >54 °C excluded as instrumental anomaly |

**Table S2.** Representative frequency-dependent rheological descriptors.

| Condition | f (Hz) | G′ (Pa) | G″ (Pa) | tanδ | \|η*\| (Pa s) |
|---|---|---|---|---|---|
| 5 mg mL$^{-1}$ LiCl | 1 | 2.11 ± 0.69 | 1.74 ± 0.42 | 0.847 ± 0.076 | 0.44 ± 0.13 |
| 5 mg mL$^{-1}$ NaCl | 1 | 1.17 ± 0.98 | 1.92 ± 1.16 | 1.958 ± 0.424 | 0.36 ± 0.24 |
| 10 mg mL$^{-1}$ LiCl | 1 | 2.673 ± 0.84 | 5.651 ± 3.30 | 2.114 ± 0.135 | 0.99 ± 0.05 |
| 10 mg mL$^{-1}$ NaCl | 1 | 22.05 ± 11.31 | 14.18 ± 5.36 | 0.724 ± 0.292 | 4.24 ± 1.83 |
| 15 mg mL$^{-1}$ LiCl | 1 | 34.95 ± 18.01 | 35.42 ± 10.95 | 1.133 ± 0.270 | 7.96 ± 3.24 |
| 15 mg mL$^{-1}$ NaCl | 1 | 108.1 ± 19.3 | 56.49 ± 4.27 | 0.531 ± 0.059 | 19.4 ± 3.0 |

**Table S3.** Selected apparent concentration-scaling and thermorheological descriptors.

| Descriptor | NaCl | LiCl | Interpretation |
|---|---|---|---|
| \|η*\| concentration exponent at ~25 °C, 1 Hz | 3.38 | 2.47 | Stronger concentration reinforcement in NaCl |
| G′ concentration exponent at ~25 °C, 1 Hz | 3.92 | 2.31 | More coherent elastic reinforcement in NaCl |
| Strongest TTS-compatible formulation | 15 mg mL$^{-1}$ | 15 mg mL$^{-1}$ (partial) | Compatibility depends on formulation |

| Descriptor | NaCl | LiCl | Interpretation |
|---|---|---|---|
| TTS apparent Ea (kJ mol−1) | 39.7 | 45.2 | Retained/compatible windows only |

**Table S4.** Global structural and ion-coordination descriptors from MD simulations. Values are mean ± SD from three independent trajectories.

| System | Max. cluster | Aggregated fraction | Rg (Å) | SASA (Å²) | Ion-O CN | Lifetime (ns) |
|---|---|---|---|---|---|---|
| NaCl 298 K | 2.257 ± 0.013 | 0.332 ± 0.008 | 51.779 ± 0.069 | 18321.14 ± 10.13 | 0.049 ± 0.013 | 2.144 ± 1.133 |
| NaCl 333 K | 2.307 ± 0.049 | 0.370 ± 0.009 | 50.660 ± 0.090 | 18315.71 ± 13.10 | 0.033 ± 0.011 | 1.427 ± 1.186 |
| LiCl 298 K | 1.693 ± 0.029 | 0.187 ± 0.010 | 51.483 ± 0.011 | 18639.69 ± 15.10 | 0.202 ± 0.009 | 19.007 ± 1.098 |
| LiCl 333 K | 1.634 ± 0.035 | 0.168 ± 0.016 | 53.345 ± 0.098 | 18625.20 ± 16.10 | 0.254 ± 0.010 | 13.607 ± 1.506 |

**Table S5.** RDF and first-shell coordination descriptors.

| System | Pair | Peak r (Å) | Peak g(r) | First minimum (Å) | Coordination number |
|---|---|---|---|---|---|
| NaCl 298 K | $Na^+$-polymer O | 2.55 | 6.970 | 3.25 | 0.0496 |
| NaCl 333 K | $Na^+$-polymer O | 2.45 | 4.655 | 3.25 | 0.0334 |
| LiCl 298 K | $Li^+$-polymer O | 2.05 | 49.490 | 2.85 | 0.2024 |
| LiCl 333 K | $Li^+$-polymer O | 2.05 | 61.114 | 3.05 | 0.2547 |

**Table S6.** Interchain ion-bridge descriptors.

| System | Bridge events | Average lifetime (ns) | Mean bridge CN | Cations | Events/100 cations/100 ns | Interpretation |
|---|---|---|---|---|---|---|
| NaCl 298 K | 3 ± 0.09 | 1.00 ± 0.06 | 3.67 ± 0.012 | 162 | 1.85 | Sparse transient events |
| NaCl 333 K | 5 ± 0.11 | 1.00 ± 0.08 | 3.40 ± 0.023 | 154 | 3.25 | Higher event frequency after heating |

| System | Bridge events | Average lifetime (ns) | Mean bridge CN | Cations | Events/100 cations/100 ns | Interpretation |
|---|---|---|---|---|---|---|
| LiCl 298 K | NA | NA | NA | 279 | 0.0 | No event under the applied criteria |
| LiCl 333 K | 9 ± 0.21 | 1.11 ± 0.02 | 3.00 ± 0.19 | 279 | 3.23 | Frequent but short-lived encounters |
